# Reduced variability of bursting activity during working memory

**DOI:** 10.1101/2022.02.18.481088

**Authors:** Mikael Lundqvist, Jonas Rose, Melissa Warden, Tim Buschman, Pawel Herman, Earl Miller

**Affiliations:** Department of Psychology, Department of clinical neuroscience, Karolinska Institute, Solna, Sweden; The Picower Institute for Learning and Memory, Department of Brain and Cognitive Sciences, Massachusetts Institute of Technology, 43 Vassar Street, Cambridge, MA, 02139, USA; Faculty of Psychology, Neural Basis of Learning, Ruhr University Bochum, 44801, Bochum, Germany; Department of Neurobiology and Behavior, Cornell University, Ithaca, NY, 14853, USA; Princeton Neuroscience Institute, Princeton University, Washington Rd., Princeton, NJ 08540, USA; Department of Computational Science and Technology, School of Electrical Engineering and Computer Science and Digital Futures, KTH Royal Institute of Technology, Stockholm, 100 44, Sweden

## Abstract

Working memories have long been thought to be maintained by persistent spiking. However, mounting evidence from multiple-electrode recording (and single-trial analyses) shows that the underlying spiking is better characterized by intermittent bursts of activity. A counterargument suggested this intermittent activity is at odds with observations that spike-time variability reduces during task performance. However, this counterargument rests on assumptions, such as randomness in the timing of the bursts, that may not be correct. Thus, we analyzed spiking and LFPs from the prefrontal cortex (PFC) of monkeys to determine if task-related reductions in variability can co-exist with intermittent spiking. We found that it does because both spiking and associated gamma bursts were task-modulated, not random. In fact, the task-related reduction in spike variability could be explained by a related reduction in gamma burst variability. Our results provide further support for the intermittent activity models of working memory as well as novel mechanistic insights into how spike variability is reduced during cognitive tasks.

## Introduction

Working memory (WM), the holding of information “online” in a form available for processing, is central in human cognition. It has long been thought that WMs are carried by persistent spiking of neurons that maintain “attractor states” (Fuster and Alexander, 1971; Funahashi et al., 1989; Goldman-Rakic, 1995; Amit and Brunel, 1997). But recent evidence suggests that WM-related activity is more dynamic (Wolff et al., 2017; Lundqvist et al., 2016; 2018; Rose et al., 2016; Sprague et al., 2016; Hussar and Pasternak, 2012; Shafi et al., 2006). Advances in multiple-electrode technology have allowed sampling of larger populations of neurons as well as a closer examination of their activity in real time. This revealed that the spiking activity of many neurons does not seem persistent. Instead, it is organized in intermittent bursts, especially when activity is examined on individual trials (Lundqvist et al., 2016; 2018; 2020; Bastos et al., 2018). Because burst times vary from trial to trial, averaging across trials can create the illusion of persistent spiking when the underlying activity is intermittent (Lundqvist et al, 2016). Further, the intermittent spiking is not just a property of individual neurons. The bursts of spiking are associated with bursts of gamma-band power (Lundqvist et al., 2016; 2018). This suggests that the intermittent bursting is coordinated in local networks. Thus, persistence cannot be obtained by pooling local networks of neurons. This provides support to a class of synaptic attractor models in which short-term plasticity helps maintain attractor states in between bouts of intermittent spiking (Wang et al. 2006; Lundqvist et al., 2011; Mongillo et al., 2008; Fiebig and Lansner, 2017). This is consistent with observations from EEG and FMRI studies that, for extended periods of time, information held in working memory cannot be decoded from global activity. However, when the cortex is “pinged” by a task-irrelevant stimulus or by transcranial magnetic stimulation, the network “rings” back with the information held in working memory (Rose et al., 2016; Sprague et al., 2016; Wolff et al., 2017).

However, observations that cortical spiking becomes less variable during performance of various tasks (Churchland et al., 2010; Cohen and Maunsell, 2009; Ponce-Alvarez, et al., 2013; Hussar and Pasternak, 2010; Ito et al, 2020) have led to a counterargument to intermittent spiking. For example, a recent modelling study argued against the synaptic attractor type models (and in favor of persistent spiking alone) by suggesting that intermittent bursting should cause an increase, not a decrease, in across-trial variability of spiking (measured by Fano Factor, FF)(Li et al., 2021). These arguments rest on several assumptions that may not be consistent with experimental observations. Most importantly, the model assumed that intermittent bursts of spiking occurred at random times during a WM task. However, experimental observations show that WM-related bursts of activity are not random, they wax and wane with different trial events (Lundqvist et al, 2016, 2018). This raises the question of whether this task-related modulation is enough to explain the task-related reduction in spike variability that is often observed in cortex.

To address this, we analyzed actual spiking and LFPs from the prefrontal cortex of non-human primates (NHPs) performing two working memory tasks. We previously reported intermittent bursts of spiking and associated bursts of gamma power during performance of the tasks (Lundqvist et al., 2016; 2018). Here, we report that variability of spiking (FF) decreased, not increased, during task performance. In fact, the largest task-related reduction in variability was found at recording sites where the spike rates and burst rates increased the most. We also gained insight into why variability was reduced. We synthesized spikes based on the gamma bursts found in the LFPs. This revealed that the reduction in spiking variability was directly related to reductions in gamma burst variability. These results provide further support for “activity-silent” synaptic attractor type models of working memory (Mongillo et al., 2008; Sandberg et al., 2003; Lundqvist et al., 2011; Fiebig and Lansner, 2017; Stokes, 2015; Kozachkov et al., 2022).

## Results

### Increased bursting co-exists with reduced spike variability

We analyzed LFP power and spiking from multiple single neurons recorded from the lateral prefrontal cortex of monkeys performing two different WM tasks. From these data, we previously reported task-related intermittent spiking and associated bursts of gamma power (Lundqvist et al., 2016; 2018). We further demonstrated that spikes associated with the gamma bursts carried more WM information than spiking without gamma bursts. Here, we examine whether the variability of this activity increases or decreases during task performance.

In both tasks, monkeys had to hold 2-3 stimuli in WM. In Task 1, the animals had to remember the color and location of 3 squares presented in their visual periphery (Lundqvist et al., 2016; Figure 1). In Task 2 – the animals remembered the identity and order of two foveally presented pictures (Warden and Miller, 2007; 2010; Supplemental Figure 1). Here, we present results from Task 1 in the main text. Results from Task 2 were used to confirm that the results generalized across tasks and experiments. Those results are presented in the Supplemental Materials.

**Figure 1.**
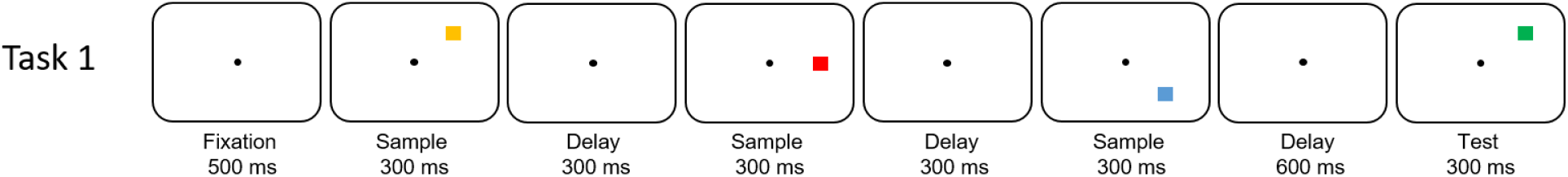
Sequential working memory change detection task. The monkeys were presented with a sequence of 3 colored squares in 3 distinct positions. After a brief delay, the monkeys were exposed to a new sequence of up to 3 squares. They reported with a saccade to the first square that had changed color between sample and test sequences (one of the squares always changed, in the above case the first).

Spiking and gamma bust rates were modulated by the task. Figure 2A shows the average spike rate as a function of time during the trial of Task 1. The gray shaded areas show the time of presentation of three to-be-remembered stimuli. As previously demonstrated, spike rate was high during the pre-stimulus baseline and dropped before presentation of the first stimulus. Presentation of each stimulus caused a transient increase in spiking. There was a ramping up of spiking over the memory delay. Figure 2B shows that gamma burst rate follows a similar profile (Supplemental Figure 2A, B shows similar results for Task 2).

**Figure 2.**
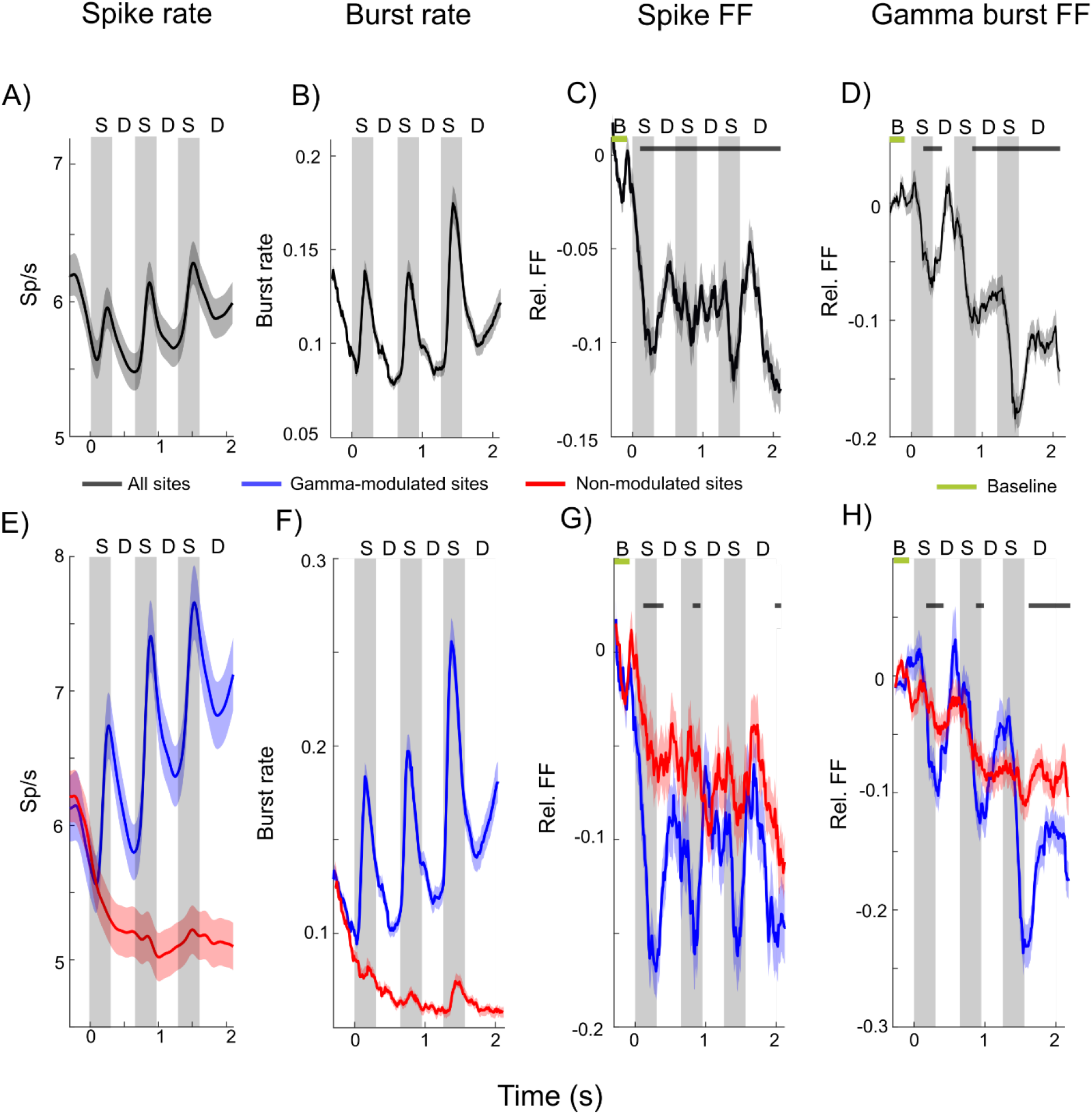
Spike and burst variability in Task 1. A) Spike rates averaged across trials and all neurons (n=495). Gray shaded regions indicate sample presentations (S; the first sample is presented at time=0). The three delays are marked by “D”. Error bars indicate standard error of the mean in all panels. B) Gamma burst rates averaged across trials for all recording sites (n=319. C) Relative FF in spike times, i.e. normalized by the pre-stimulus baseline (B; 300 to 100 ms prior to first sample onset) average. D) Relative FF in burst times (normalization as in C). Black bars indicate time periods when post stimulus FF deviated significantly from the pre-stimulus baseline (marked by the green bar, p<0.0001, permutation based cluster statistics (Maris and Oostenveld, 2007)). E-H) same as A-D) but separately for sites that increased (blue; n=145 sites and n=214 neurons) gamma bursting at sample onset, and sites that did not (red; n=174 sites and n=281 neurons). Black bars indicate time periods when blue and red curves are significantly different (p<0.03 permutation based cluster statistics).

This modulation of spiking and gamma bursting by the task resulted in a decrease in across-trial variability. This can be seen in Figures 2C and 2D, respectively, which plot the change in Fano Factor (FF) relative to the pre-stimulus baseline over the trial duration. The FF was high during the baseline and decreased during task performance. Importantly, both FFs show opposite trends to spike and gamma burst rates (c.f. Figures 2A-B). The FFs decrease during stimulus presentations, increase between stimulus presentations and decrease towards the end of the delay. Notably, both the spike and gamma burst FFs were lower during the memory delay than during the pre-stimulus baseline.

Our prior work showed a relationship between gamma bursts and WM information in spiking. We found that stimulus-driven gamma bursting was limited to a subset of recording sites (145/319=45% in Task 1, 151/199=73% in Task 2; sites with significant increase in bursts during stimulus presentation relative to the pre-stimulus period, p=0.05, see Lundqvist et al, 2016 for Task 1, Lundqvist et al., 2018 for Task 2). We referred to these sites as “gamma-modulated sites” while the sites without stimulus-induced gamma bursting were referred to as “non-modulated sites” (Lundqvist et al, 2016, 2018). We then demonstrated that spiking at gamma-modulated sites exclusively carried information about the stimuli held in WM.

Both gamma-modulated and non-modulated sites showed a reduction in gamma-burst variability during the task. Figures 2E and 2F show the difference in spike rate (Figure 2E) and gamma burst rate (Figure 2F) between gamma-modulated and non-modulated sites. Much like the average across all sites (Figure 2B), gamma burst rate at gamma-modulated sites increased during stimulus presentation and ramped up over the memory delay (Figure 2F, blue lines). As expected, there was less gamma bursting and no stimulus dependent modulation at non-modulated sites (Figure 2F, red lines), and the spike rates were lower at non-modulated sites (Figure 2E).

We also found that the spiking FF showed a greater reduction at the gamma-modulated sites (Figure 2G, blue lines) than non-modulated sites (Figure 2G, red lines). Gamma bursting at both types of sites showed a reduction in FF as the trial progressed relative to the baseline but this effect was more pronounced at the gamma modulated sites (Figure 2H). During stimulus presentations, when both gamma burst and spike rates increased at the gamma-modulated sites, there were sharp drops in the gamma burst FF at these sites.

These analyses indicate that task modulation of spike activity and gamma bursts does result in task-related reduction in neural variability. The examination of gamma-modulated vs non-modulated sites further indicates that the more task-modulation of bursting, the greater the reduction in both spike and gamma burst variability. The gamma bursts are thought to reflect coordinated spiking in local networks. This raises the question of whether the task-modulation of the gamma bursts can, in turn, explain the task-related reduction in spike variability. We address this below.

### Reductions of gamma burst variability can explain the reduction in spike variability

To further examine the relationship between spiking and gamma burst variability, we used gamma burst events to construct synthetic spike trains. In particular, inside every gamma burst identified (see Methods and Lundqvist et al., 2016) at every recording site on each trial we generated an artificial Poisson spike train (see Methods). We systematically varied the spike rates inside bursts over a wide range and quantified the resulting synthetic spike variability across trials using FF. The analysis revealed that task-related modulations of the gamma bursts reduced the FF for the artificial spikes just as it did for actual recorded spikes (Figure 3; Supplemental Figure 3). In fact, this effect was observed across a wide range of spike rates within the bursts, although it was more pronounced for the higher spiking rates. The effect gradually decreased as spike rates inside bursts were reduced to 10 sp/s (corresponding to the average of 0.54 sp/burst given the average burst duration of 54 ms in the data. Thus bursting doesn’t have to be apparent in single neuron activity necessarily). In addition, we added background spiking activity outside bursts (at 1 sp/s). This resulted in a weakening of the stimulus-driven reduction in spike time variability, especially for low within-burst firing rates (Figure 3B). This suggests that most spiking has to occur during the bursts for the gamma burst variability to drive the observed drop in spike variability. Thus in the burst-driven spiking regime, the reduction in spiking variability during the task can be explained by the reduction of gamma burst variability.

**Figure 3.**
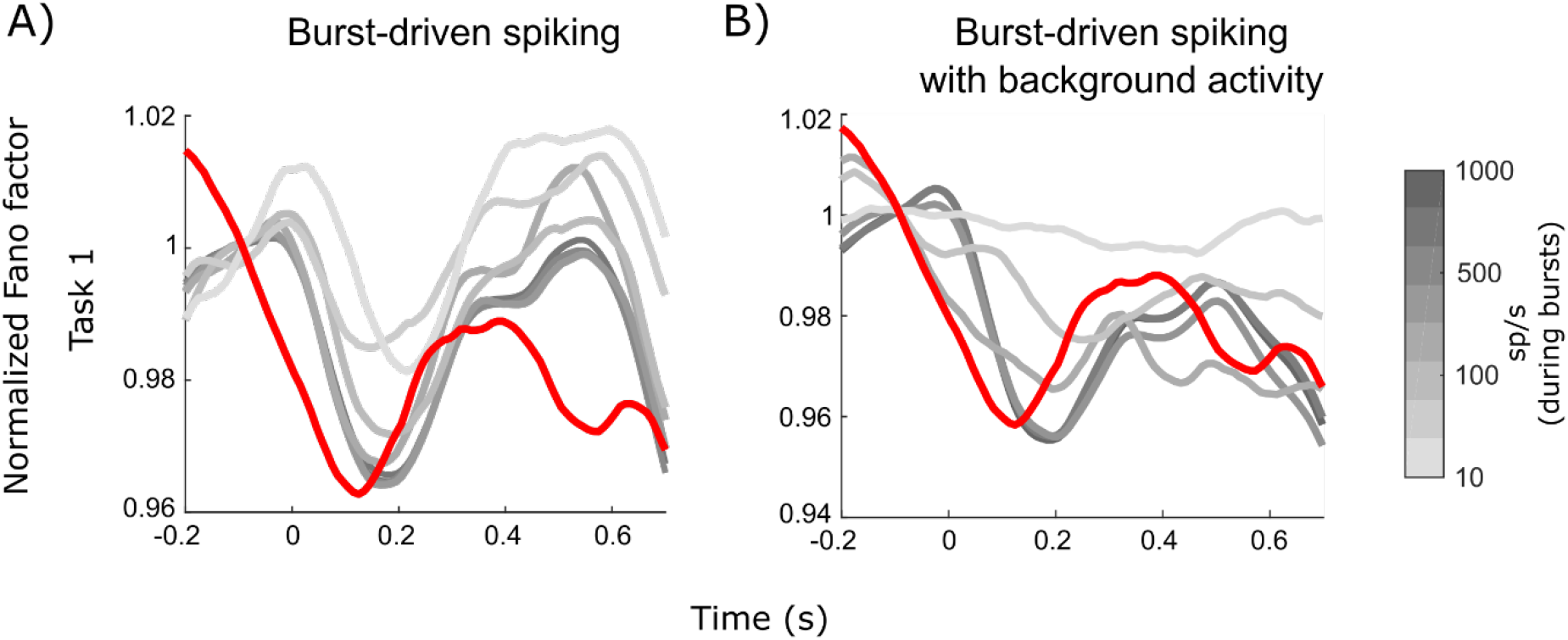
Variability of synthetic spike trains generated based on neurophysiological gamma burst times in Task 1. Spiking rates within the bursts were varied from 10 to 1000 sp/s (black to light gray). A) Plotted are the mean (averaged across all the recording sites) FFs normalized divisively by the pre-stimulus average, for a range of simulated spiking rates without (A) and with (B) an additional synthetic background activity at the level of 1sp/s. Red line corresponds to the original normalized spike FF for the recorded spiking. We focused on the first sample presentation to exploit more trials with identical conditions (for each sample presentation the conditions gradually branched out so the proportional share of particular stimulus combinations decreased).

## Discussion

We found that the task-related modulation of bursts of spiking and gamma power during a WM task resulted in the cross-trial reduction in the variability of neural activity. Further, we found a direct relationship between the reduction of the variability of gamma bursting and the reduction of spiking variability. They co-occurred both in time and space. The reduced spike variability was most pronounced during task events and at recording sites where the reduction in gamma burst FF was large. These were the same sites where gamma bursting and spiking were most strongly task-modulated. Further, reductions in activity variability were seen at recording sites with increased and decreased gamma bursting. Thus, the reduced burst variability was not due to increased bursting per se. This can explain why the reduction in spiking variability is largely decoupled from spike rates (Hussar and Pasternak, 2010). It was the increase in the structure of bursting activity via modulation by the task that produced the reduction in variability. We investigated this by synthesizing spike trains from recorded gamma burst patterns and found that the pattern of bursts could explain reduced spike variability for a wide range of synthetic spike rates.

These results provide further support for working memory models in which information is held by a combination of spiking (during bursts) and short-term synaptic plasticity (between bursts) rather than persistent spiking alone (Sandberg et al., 2003; Mongillo et al., 2008; Lundqvist et al., 2011; Fiebig and Lansner, 2017; Kozachkov et al., 2022). In these “synaptic attractor” models bursts of spiking induces temporary synaptic imprints that maintain a trace of the attractor state in between spiking. Further, recent evidence has shown that the short-term synaptic plasticity endows additional benefits beyond memory maintenance. For instance, it helps make recurrent networks more stable and more robust to perturbations and synaptic loss (Kozachkov et al., 2022; Wu and Zenke, 2021).

A recent computation modeling study argued that intermittent bursts of activity is not compatible with task-related reductions in spiking variability (Li et al, 2021). However, this model used several problematic assumptions that were not supported by experimental observation. First, the Li et al. (2021) model assumed intermittent bursting occurred at random times during tasks. This is not correct. Experimental observations have shown that bursting is task-modulated, waxing and waning with different trial events (Lundqvist et al, 2016, 2018). Second, their model compared synthetic task-related activity to a pre-task baseline during which there were no random bursts of (synthetic) spikes. There was only steady background spiking. This is also incorrect. There is random bursting during pre-task baselines (Lundqvist et al., 2016; 2018). Thus, using an artificial steady-state (non-bursting) baseline meant that they made comparisons to a baseline with a low-level of variability not actually seen in cortex. Third, in their model the task-related reductions in neural variability were highly dependent on model assumptions. The model could show an increase vs decrease in variability depending on whether spike rates were increased by increasing the number vs duration of the bursts (Equation 9 in Li et al., 2021). They chose to increase the duration of the bursts. However, experimental observations have shown that burst rates increase but their duration remains constant (Lundqvist et al., 2016). We avoided problems with model assumptions by not making any assumptions. Instead, we used activity recorded from the PFC of non-human primates.

With our ability to record and analyze ever larger quantities of simultaneous neuron activity, there has been increasing evidence for discrete, packet-based activity in cortex (Lakatos et al., 2008; Buschman and Miller, 2010; Schroeder et al, 2010; Engel et al., 2016; Luczak et al., 2015; Helfrich et al, 2018; VanRullen, 2018; Stringer et al., 2019; Lundqvist et al., 2016; Lundqvist and Wutz, 2021). In this view, underlying cortical processing is discrete events that form packets (bursts) of processed information. It allows time multiplexing such as models of multi-item working memory where each item takes turn being active and silent (Lisman and Idiart, Lundqvist et al., 2011). It would also facilitate inter-areal communication as packets of finite and standardized size are sent and received (Womelsdorf and Fries, 2006; Buschman and Miller, 2010; Buffalo, 2011; Bastos et al, 2015). Slow oscillations in the theta and sometimes alpha range have long been proposed to aid such coordinated inter-areal activities (Landau et al., 2015; Cavanagh et al., 2009; Sauseng et al., 2004). These oscillations occur at a similar time scale as the spike bursts and are related (Canolty et al., 2006). Here we demonstrate that this framework is consistent with task-related reductions in neuron variability that is common across cortex.

## Supplemental Figures

**Supplemental Figure 1.**
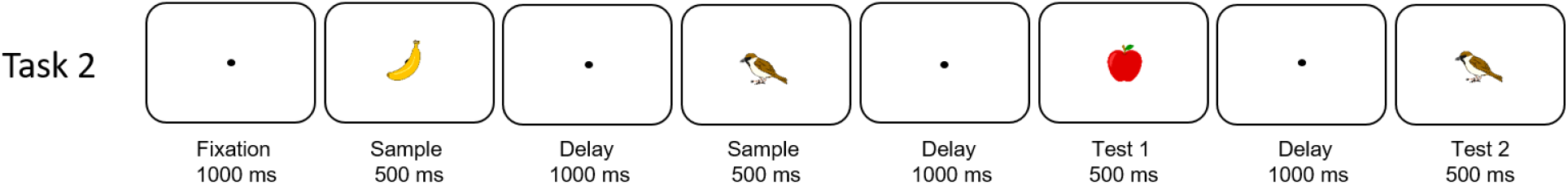
Structure of Task 2. The monkeys were presented with a sequence of 2 objects foveally. After a brief delay, the monkeys were exposed to a new sequence of 2 objects. They reported by lifting a bar if the test sequence did not match the sample sequence.

**Supplemental Figure 2.**
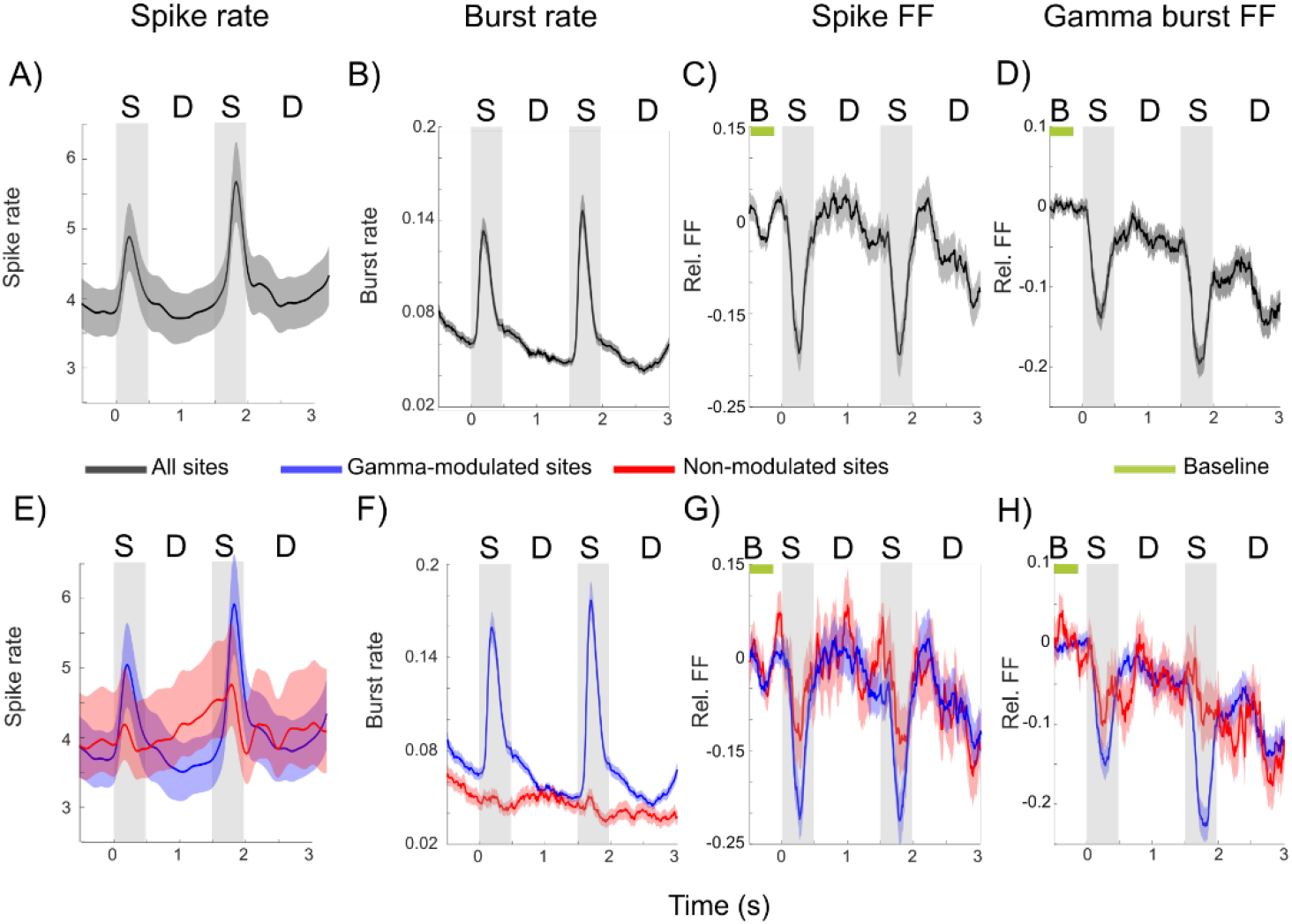
Spike and burst variability in Task 2. A) Spike rates averaged across trials and all neurons (n=283). Gray shaded regions indicate sample presentations (S; the first sample is presented at time=0). The two delays are marked by D. Error bars indicate standard error of mean in all panels. B) Gamma burst rates averaged across trials for all recording sites. C) FF in spike times normalized by the pre-stimulus baseline (B; 500 to 100 ms prior to first sample onset) average. D) FF in burst times normalized by the pre-stimulus (500 to 100 ms prior to first sample onset) average. E-H) same as A-D) but separately for sites that increased (blue) gamma bursting at sample onset, and sites that did not (red).

**Supplemental Figure 3.**
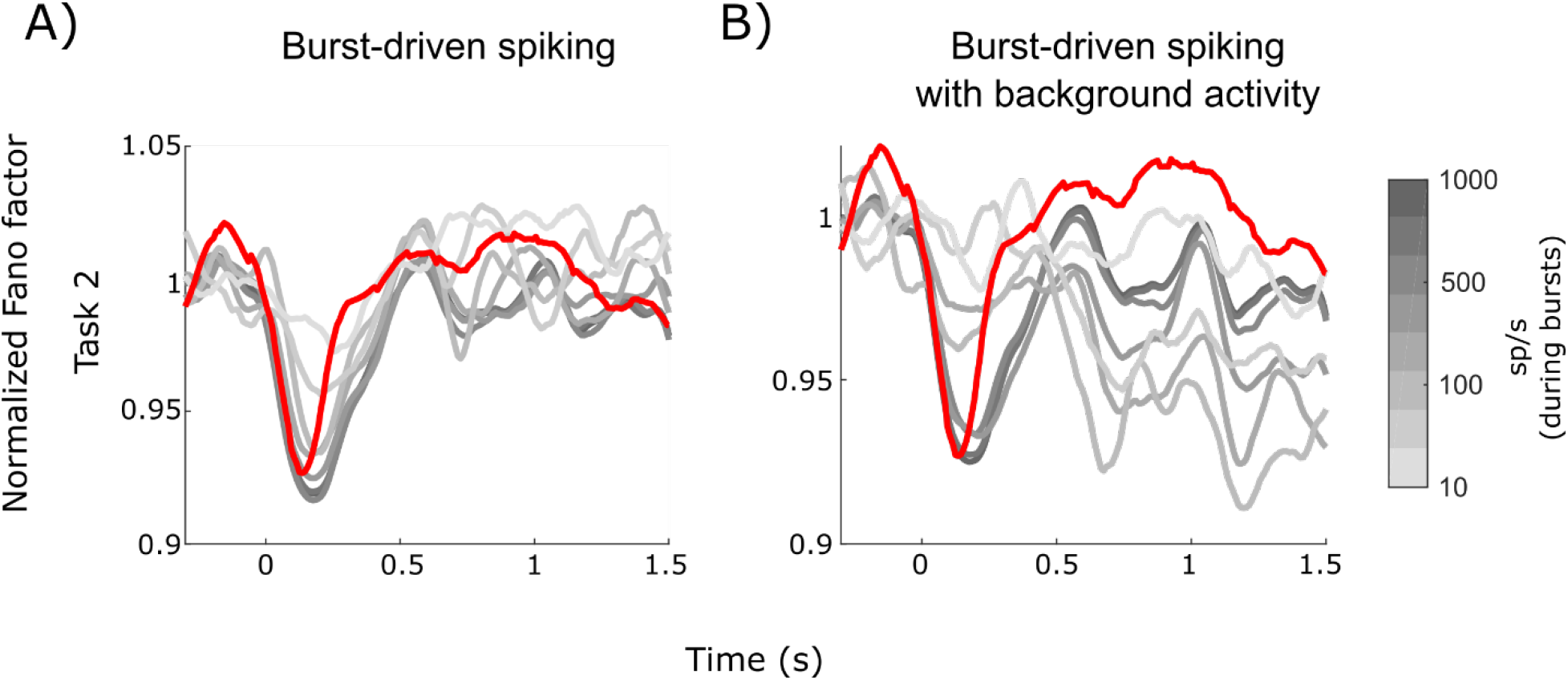
FF for synthetic spike trains generated based on neurophysiological gamma burst times in Task 2. Spike rates within the bursts were varied from 10 to 1000 sp/s (black to light gray). A) Plotted are the normalized FFs (divided by the pre-stimulus average FF) for different spike rates, averaged across all recording sites. Red line plots the original spike FF, normalized in the same way. B) same as A) but with background Poisson activity at 1 sp/s added.

## Materials and Methods

### Data

We analyzed data from prior studies (Warden and Miller, 2007; 2010; Lundqvist et al., 2016; 2018). This data consisted of two data sets with acute electrode recordings from PFC while monkeys performed two distinct WM tasks (Task 1, Task2, Figure 1, Suppl Figure 1). Task 1 corresponds to 3 item-trials in Lundqvist et al., 2016. We focused on 3-item trials as it allowed us to have a larger number of repeated trials. Task 2 corresponds to the recognition task in (Warden and Miller, 2007; 2010), which is the same data analyzed in (Lundqvist et al., 2018). The animals received postoperative antibiotics and analgesics and were always handled in accord with the National Institutes of Health guidelines, and all procedures were approved by the Massachusetts Institute of Technology Committee on Animal Care.

The LFPs were recorded at 30 kHz in Task 1 and 1 kHz in Task 2. The data from Task 1 was down sampled to 1 kHz before further analysis. We kept all electrode contacts with at least one associated isolated neuron. As a result, we obtained 495 units corresponding to 319 electrodes in Task 1 and X units on Y electrodes in Task 2. Only correct trials were kept for further analysis (on average, 255+-72 trials in Task 1 and 332+-80 trials in Task 2).

Gamma modulated sites were chosen based on activity during sample presentations. To this end, we identified sites with the average gamma burst rate over any of the sample presentations significantly exceeding the threshold of 0.03. The rest of sites were considered as non-modulated.

### Burst analysis

We relied on gamma bursts times extracted in the two prior studies (Lundqvist et al., 2016; 2018). In short, we first extracted the gamma-band power in 3 distinct bands (40-65, 55-90 and 70-100 Hz). For this purpose we adopted a multi-taper approach with frequency-dependent window lengths corresponding to 6-8 oscillatory cycles and frequency smoothing corresponding to 0.2-0.3 of the central freq, *f*_0_, i.e. *f*_0_±0.2*f*_0_, where *f*_0_ were sampled with the resolution of 1 Hz (this configuration implies that 2-3 tapers were used). We then extracted an estimate of the band power envelope by averaging spectral components within each band with the temporal resolution of 1 ms. Next, we thresholded the envelopes based on their mean and standard deviation during fixation in pre-stimulus conditions (using the last 300 ms prior to stimulus onset in Task 1, last 500 ms prior to stimulus onset in Task 2).

A gamma burst was defined as a time interval when the envelope exceeded the mean by two standard deviations for at least the duration of 3 oscillatory cycles (using the center of each gamma band to define the length of an oscillatory cycle). In line with our previous work (Lundqvist et al., 2016; 2018) we also used a trial-average measure of burst rates for each spectral band. It corresponds to the chance of a burst occurrence on an individual electrode at a particular time in the trial. In this work, we used the union of bursts across the three gamma bands and, accordingly, we relied on the average burst rate across those identified for the three gamma bands.

### Fano factor (FF)

We estimated the spiking Fano factor (FF) for each unit in a two-step process. First, we calculated the number of spikes, *s*, within a sliding window of 150 ms and calculated its mean, *μ_s_,* and variance, *σ_s_*^2^, across trials to estimate FF = *σ_s_*^2^/ *μ_s_*. This resulted in a raw FF time series of 1 ms resolution. Further, to facilitate comparisons we show in Figure 2 and Supplemental Figure 2 the relative FF, which is obtained by removing the mean FF over the pre-stimulus baseline from the raw FF.

The FF for bursts was estimated using a binary time series for each site and trial with 0 corresponding to the time bin outside a burst event and 1 – inside a burst event. The resulting time series was then subject to the same treatment as spike time series.

### Spike train analysis and generation

To estimate spiking rates we convolved spike trains with a 50-ms-wide Gaussian kernel. We constructed synthetic spike trains based on the recorded burst timings. Within each burst interval we generated spikes from a Poisson process at the desired rate ranging between 10 and 1000 sp/s. In another manipulation (Figure 3B) we also generated background spike activity with the constant rate of 1 sp/s.

## Acknowledgements

This work was supported by ONR MURI N00014-16-1-2832, ERC starting grant 949131, NIMH R37MH087027, and The JPB Foundation, by the Swedish Research Council (Vetenskapsrådet) 2018-05360, 2018-04197 and 2016-05871.

## References

Amit, D. J., & Brunel, N. (1997). Model of global spontaneous activity and local structured activity during delay periods in the cerebral cortex. Cerebral cortex (New York, NY: 1991), 7(3), 237–252.

Bastos, A. M., Vezoli, J., & Fries, P. (2015). Communication through coherence with inter-areal delays. Current opinion in neurobiology, 31, 173–180.

Bastos, A. M., Loonis, R., Kornblith, S., Lundqvist, M., & Miller, E. K. (2018). Laminar recordings in frontal cortex suggest distinct layers for maintenance and control of working memory. Proceedings of the National Academy of Sciences, 115(5), 1117–1122.

Buffalo, E. A., Fries, P., Landman, R., Buschman, T. J., & Desimone, R. (2011). Laminar differences in gamma and alpha coherence in the ventral stream. Proceedings of the National Academy of Sciences, 108(27), 11262–11267.

Buschman, T.J. and Miller, E.K. (2010) Shifting the Spotlight of Attention: Evidence for Discrete Computations in Cognition. Frontiers in Human Neuroscience. 4(194): 1–9

Canolty, R. T., Edwards, E., Dalal, S. S., Soltani, M., Nagarajan, S. S., Kirsch, H. E., … & Knight, R. T. (2006). High gamma power is phase-locked to theta oscillations in human neocortex. science, 313(5793), 1626–1628.

Cavanagh, J. F., Cohen, M. X., & Allen, J. J. (2009). Prelude to and resolution of an error: EEG phase synchrony reveals cognitive control dynamics during action monitoring. Journal of Neuroscience, 29(1), 98–105.

Churchland, M. M., Byron, M. Y., Cunningham, J. P., Sugrue, L. P., Cohen, M. R., Corrado, G. S., … & Shenoy, K. V. (2010). Stimulus onset quenches neural variability: a widespread cortical phenomenon. Nature neuroscience, 13(3), 369–378.

Cohen, M. R., & Maunsell, J. H. (2009). Attention improves performance primarily by reducing interneuronal correlations. Nature neuroscience, 12(12), 1594–1600.

Engel, T. A., Steinmetz, N. A., Gieselmann, M. A., Thiele, A., Moore, T., & Boahen, K. (2016). Selective modulation of cortical state during spatial attention. Science, 354(6316), 1140–1144. https://doi.org/10.1126/science.aag1420

Goldman-Rakic, P. S. (1995). Cellular basis of working memory. Neuron, 14(3), 477–485.

Helfrich, R. F., Fiebelkorn, I. C., Szczepanski, S. M., Lin, J. J., Parvizi, J., Knight, R. T., & Kastner, S. (2018). Neural mechanisms of sustained attention are rhythmic. Neuron, 99(4), 854–865.

Hussar, C., & Pasternak, T. (2010). Trial-to-trial variability of the prefrontal neurons reveals the nature of their engagement in a motion discrimination task. Proceedings of the National Academy of Sciences, 107(50), 21842–21847.

Hussar, C. R., & Pasternak, T. (2012). Memory-guided sensory comparisons in the prefrontal cortex: contribution of putative pyramidal cells and interneurons. Journal of Neuroscience, 32(8), 2747–2761.

Fiebig, F., & Lansner, A. (2017). A spiking working memory model based on Hebbian short-term potentiation. Journal of Neuroscience, 37(1), 83–96.

Funahashi S, Bruce CJ, Goldman-Rakic PS (1989) Mnemonic coding of visual space in the monkey’s dorsolateral prefrontal cortex. J Neurophysiol 61:331–349.

Fuster JM, Alexander GE (1971) Neuron activity related to short-term memory. Science 173:652–654.

Kozachkov, L., Tauber, J., Lundqvist, M., Brincat, S. L., Slotine, J. J., & Miller, E. K. (2022). Robust Working Memory through Short-Term Synaptic Plasticity. bioRxiv.

Ito, T., Brincat, S. L., Siegel, M., Mill, R. D., He, B. J., Miller, E. K., … & Cole, M. W. (2020). Task-evoked activity quenches neural correlations and variability across cortical areas. PLoS computational biology, 16(8), e1007983.

Landau, A. N., Schreyer, H. M., Van Pelt, S., & Fries, P. (2015). Distributed attention is implemented through theta-rhythmic gamma modulation. Current Biology, 25(17), 2332–2337.

Lakatos, P., Karmos, G., Mehta, A. D., Ulbert, I., & Schroeder, C. E. (2008). Entrainment of neuronal oscillations as a mechanism of attentional selection. science, 320(5872), 110–113.

Li, D., Constantinidis, C., & Murray, J. D. (2021). Trial-to-trial variability of spiking delay activity in prefrontal cortex constrains burst-coding models of working memory.

Lisman, J. E., & Idiart, M. A. (1995). Storage of 7±2 short-term memories in oscillatory subcycles. Science, 267(5203), 1512–1515.

Luczak, A., McNaughton, B. L., & Harris, K. D. (2015). Packet-based communication in the cortex. Nature Reviews Neuroscience, 16(12), 745–755.

Lundqvist, M., Herman, P., & Lansner, A. (2011). Theta and gamma power increases and alpha/beta power decreases with memory load in an attractor network model. Journal of cognitive neuroscience, 23(10), 3008–3020.

Lundqvist, M., Rose, J., Herman, P., Brincat, S. L., Buschman, T. J., & Miller, E. K. (2016). Gamma and beta bursts underlie working memory. Neuron, 90(1), 152–164.

Lundqvist, M., Herman, P., Warden, M. R., Brincat, S. L., & Miller, E. K. (2018). Gamma and beta bursts during working memory readout suggest roles in its volitional control. Nature communications, 9(1), 1–12.

Lundqvist, M., Bastos, A. M., & Miller, E. K. (2020). Preservation and changes in oscillatory dynamics across the cortical hierarchy. Journal of cognitive neuroscience, 32(10), 2024–2035.

Lundqvist, M., & Wutz, A. (2021). New methods for oscillation analyses push new theories of discrete cognition. Psychophysiology, e13827.

Maris, E., & Oostenveld, R. (2007). Nonparametric statistical testing of EEG-and MEG-data. Journal of neuroscience methods, 164(1), 177–190.

Mongillo, G., Barak, O., & Tsodyks, M. (2008). Synaptic theory of working memory. Science, 319(5869), 1543–1546.

Ponce-Alvarez, A., Thiele, A., Albright, T. D., Stoner, G. R., & Deco, G. (2013). Stimulus-dependent variability and noise correlations in cortical MT neurons. Proceedings of the National Academy of Sciences, 110(32), 13162–13167.

Sandberg, A., Tegnér, J., & Lansner, A. (2003). A working memory model based on fast Hebbian learning. Network: Computation in Neural Systems, 14(4), 789.

Sauseng, P., Klimesch, W., Doppelmayr, M., Hanslmayr, S., Schabus, M., & Gruber, W. R. (2004). Theta coupling in the human electroencephalogram during a working memory task. Neuroscience letters, 354(2), 123–126.

Shafi, M., Zhou, Y., Quintana, J., Chow, C., Fuster, J., & Bodner, M. (2007). Variability in neuronal activity in primate cortex during working memory tasks. Neuroscience, 146(3), 1082–1108.

Schroeder, C. E., Wilson, D. A., Radman, T., Scharfman, H., & Lakatos, P. (2010). Dynamics of active sensing and perceptual selection. Current opinion in neurobiology, 20(2), 172–176.

Stokes, Mark G. “‘Activity-silent’ working memory in prefrontal cortex: a dynamic coding framework.” Trends in cognitive sciences 19.7 (2015): 394–405.

Stringer, C., Pachitariu, M., Steinmetz, N., Reddy, C. B., Carandini, M., & Harris, K. D. (2019). Spontaneous behaviors drive multidimensional, brainwide activity. Science, 364(6437). https://doi.org/10.1126/science.aav7893

Sprague, T. C., Ester, E. F., & Serences, J. T. (2016). Restoring latent visual working memory representations in human cortex. Neuron, 91(3), 694–707.

Rose, N. S., LaRocque, J. J., Riggall, A. C., Gosseries, O., Starrett, M. J., Meyering, E. E., & Postle, B. R. (2016). Reactivation of latent working memories with transcranial magnetic stimulation. Science, 354(6316), 1136–1139.

VanRullen, R. (2018). Attention cycles. Neuron, 99(4), 632–634.

Warden, M. R., & Miller, E. K. (2010). Task-dependent changes in short-term memory in the prefrontal cortex. Journal of Neuroscience, 30(47), 15801–15810.

Warden, M. R., & Miller, E. K. (2007). The representation of multiple objects in prefrontal neuronal delay activity. Cerebral Cortex, 17(suppl_1), i41–i50.

Wang, Y., Markram, H., Goodman, P. H., Berger, T. K., Ma, J., & Goldman-Rakic, P. S. (2006). Heterogeneity in the pyramidal network of the medial prefrontal cortex. Nature neuroscience, 9(4), 534–542.

Wolff, M. J., Jochim, J., Akyürek, E. G., & Stokes, M. G. (2017). Dynamic hidden states underlying working-memory-guided behavior. Nature neuroscience, 20(6), 864–871.

Womelsdorf, T., & Fries, P. (2006). Neuronal coherence during selective attentional processing and sensory–motor integration. Journal of Physiology-Paris, 100(4), 182–193.

Wu, Y. K., & Zenke, F. (2021). Nonlinear transient amplification in recurrent neural networks with short-term plasticity. Elife, 10, e71263.

